# Pathological calcium influx through amyloid beta pores disrupts synaptic function

**DOI:** 10.1101/2025.03.27.645755

**Authors:** Temitope Adeoye, Ghanim Ullah

## Abstract

Alzheimer’s disease (AD) is characterized by profound disruption of synaptic function, with mounting evidence suggesting that amyloid-β (Aβ) oligomers disrupt calcium (Ca^2+^) homeostasis through membrane pore formation. While these pores are known to alter intracellular Ca^2+^ dynamics, their immediate impact on synaptic transmission and potential interaction with Familial AD (FAD)-associated endoplasmic reticulum (ER) dysfunction remains unclear. Here, we extend our previously developed model of presynaptic Ca^2+^ dynamics to examine how Aβ pore activity alters exocytosis and how such disruptions may manifest in the presence of FAD-associated ER dysfunction. Our model reveals that Aβ pores fundamentally alter both the timing and strength of neurotransmitter release. Unexpectedly, the impact of pores on synaptic function depends critically on their pattern of activity, where continuous pore activity leads to synaptic hyperactivation, while temporally brief periods of intense pore activity trigger lasting hypoactivation at short timescales. These effects manifest most strongly in synapses with low and intermediate release probabilities, highlighting the established selective vulnerability of such synaptic configurations. We find that Aβ pores and FAD-driven ER Ca²⁺ dysregulation form an integrated pathological unit through bidirectional coupling of their respective Ca²⁺ microdomains to create complex patterns of disruptions. This coupling creates feedback loops that produces an additive effect on neurotransmitter release during brief stimulations, but non-additive effects during sustained activity that promotes a shift towards asynchronous release. Surprisingly, our simulations predict that extended pore activity does not worsen indefinitely but only produces a modest additional disruption beyond initial pore formation that is likely determined by the intrinsic properties of the synapse. These findings indicate that early synaptic dysfunction in AD may arise from subtle perturbations in the temporal coordination of release rather than gross Ca^2+^ dysregulation, providing new mechanistic insights into the progressive nature of Aβ-driven synaptic failure in AD.

## Introduction

Alzheimer’s disease (AD) is characterized by extracellular deposition of senile amyloid (Aβ) plaques and tau neurofibrillary tangles which contribute to synaptic dysfunction and neuronal loss underlying cognitive decline in AD^1–4^. Specifically, the accumulated neuritic plaques trigger impaired neurotransmitter release, aberrant synaptic plasticity, and functional disruption of neuronal networks^5–8^. The precise molecular mechanisms underlying the deleterious effects of soluble Aβ oligomers on synaptic function and neurotransmission in AD remain an area of active investigation. Nevertheless, substantial evidence indicates that these peptides disrupt intracellular calcium (Ca^2+^) signaling pathways, leading to impaired Ca^2+^ homeostasis and synaptic dysfunction^9,10^. For instance, Aβ oligomers potentiate NMDA receptor-mediated Ca^2+^ influx and production of reactive oxygen species in hippocampal cultures^11^. Likewise, Ca^2+^ dyshomeostasis has been shown to precede synapse loss in human cerebral cortices treated with Aβ peptides, supporting the crucial involvement of Ca^2+^-mediated Aβ neurotoxicity *in vivo*^12^.

A key mechanism proposed to underly Aβ-mediated Ca^2+^ toxicity is the formation of Ca^2+^ channels which disrupt plasma membrane integrity and subsequently causes dysregulated Ca^2+^ influx^9,13–16^. In agreement with this idea, numerous studies utilizing lipid bilayers and Xenopus oocytes lacking native plasma membrane Ca^2+^ channels have consistently demonstrated the formation of these Ca^2+^-permeable pores by Aβ oligomers on the cell membrane^15,17–21^. Likewise, studies in organotypic hippocampal cultures have shown that direct Aβ-mediated perforation of the native neuronal membranes leads to increase in intracellular Ca^2+^ and a Ca^2+^-dependent depletion of vesicular resources^22,23^. These findings corroborate the ion-channel hypothesis of AD, which posits that soluble Aβ oligomers alter cellular ionic homeostasis and induce synaptotoxicity by forming Ca^2+^-permeable pores^16,24^.

Intracellular Ca^2+^ plays a crucial role in regulating various neuronal functions, including neurotransmitter release. The precise temporal regulation of synaptic transmission by Ca^2+^ is primarily achieved through voltage-gated Ca^2+^ channels (VGCCs), which mediate the rapid transduction of depolarization-induced Ca^2+^ transients into exocytosis^25–28^. Furthermore, Ca^2+^ influx through VGCCs is involved in a variety of physiological events underpinning protein expression, spine maintenance, and the regulation of excitability in excitatory synapses^29–33^. Since Ca^2+^ handling is critical for presynaptic function, altered Ca^2+^ homeostasis gives rise to a myriad of dysregulated neuronal processes in AD, leading to the Ca^2+^ hypothesis of AD and aging^34^. As Ca^2+^ primarily regulates the biological machinery responsible for exocytosis and short-term plasticity^35^, perturbations in Ca^2+^ handling particularly exert profound impact on synaptic transmission. These disruptions, coupled with the enhanced vulnerability of neurons to apoptotic stimuli triggered by a Ca^2+^-dependent elevation of glutamate excitotoxity, underscore the critical role of Ca^2+^ processing in mediating the deleterious effects of Aβ in affected cells^36^. Therefore, it is especially valuable to elucidate the neurotoxic nature of Aβ-induced Ca^2+^ permeable pores and their effect on Ca^2+^ signaling pathways and other presynaptic processes that are highly sensitive to pathological perturbations.

Growing evidence suggests that these pores, though atypical of single-channel behavior, exhibit increased toxicity over time^23^. Exposure of rat hippocampal neurons to Aβ results in a significant and persistent increase in intracellular Ca^2+^ levels via the formation of Aβ pores in the presynaptic neuronal membrane. This in turn causes a transient increase in transmitter release lasting up to four hours, followed by an acute decrease in excitatory miniature currents, indicative of ensuing synaptic failure^22^. Moreover, perforated patch clamp recordings and single-cell imaging revealed the temporal instability Aβ pores, which transition from Cl^-^-selective to non-selective pores, leading to a disruption of the cationic gradient and enabling entry of larger molecules^37^. These findings suggest that Aβ may initiate its own nucleation, leading to progressively larger perforations in the membrane. Furthermore, we recently demonstrated the dynamic nature of plasma membrane Aβ_42/40_ pores, which trigger Ca^2+^ toxicity that accelerates cell death. Monitoring the activity of thousands of these pores over tens of minutes revealed that not only does the number of Aβ_40_ pores increase over time, but also the Ca^2+^ toxicity of pre-existing pores continues to evolve due to increases in their open probability and Ca^2+^ permeability^38,39^. These results underscore the intricate mechanisms through which Aβ disrupts membrane integrity and the need for a comprehensive exploration of the potential link between disrupted Ca^2+^ homeostasis and impaired synaptic function in AD.

Among other mechanisms necessary for normal neuronal function, there is extensive evidence supporting a critical role of intracellular stores in maintaining precise spatiotemporal Ca^2+^ signaling events in the nerve terminal of neurons, especially at CA3 to CA1 synapses in the hippocampus^40–42^. A recent *in silico* study suggests that the presynaptic endoplasmic reticulum (ER) is critical in maintaining short-term plasticity and enabling reliable operation of synapses with low release probability^43^. These stores are susceptible to pathological perturbations in Ca^2+^ handling, which ultimately compromise synaptic transmission and plasticity^44^. Indeed, both intra- and extracellular Aβ oligomers as well as FAD-causing mutations in presenilin result in enhanced Ca^2+^ release from the ER via inositol (1,4,5)-triphosphate (IP_3_) receptors (IP_3_Rs) and/or ryanodine receptors (RyRs), contributing to aberrant plasticity. In a recent study^48^, we used a detailed biophysical model of Ca^2+^-driven exocytosis at the CA3 presynaptic terminal to study the pathological role of aberrant ER-driven neuronal Ca^2+^ handling in glutamate release and presynaptic dysfunction observed in AD. Our study demonstrated how FAD-associated ER perturbations alter presynaptic neurotransmitter release rates and synaptic plasticity and facilitation at affected synapses. While it is well established that Aβ pores and aberrant ER function can alter similar intracellular signaling pathways, the precise manner in which Ca^2+^ homeostasis is affected by the interplay between aberrant ER function, the kinetics and evolution of Ca^2+^ toxicity induced by Aβ pores, as well as their collective impact on the different aspects of neurotransmitter release, is not fully understood and challenging to experimentally assess. Of critical importance are the unanswered questions regarding how these interactions contribute to impaired synaptic plasticity and the subsequent cognitive decline observed in AD.

In the present study, we extend the same in-silico model of action potential (AP)-driven neurotransmission at the CA3 presynaptic terminal, considering the prolonged effect of Aβ pores on intracellular Ca^2+^ handling and exocytosis, including their interplay with FAD-driven ER perturbations on presynaptic integrity. Our model includes a flux representing the rate of Ca^2+^ entry through Ca^2+^ permeable pores on the membrane formed by Aβ peptides, accounting for the kinetics and evolution of the pores, as well as their co-localization with the active zone. By incorporating critical aspects of Ca^2+^ signaling, Aβ pore formation, and FAD-associated pathological Ca^2+^ release from the ER, our model robustly captures the complex interplay of key driving mechanisms for Ca^2+^ disruption in AD and their combined effect on presynaptic neurotransmitter release, synaptic plasticity, and facilitation at affected synapses. Overall, this work provides novel insights in to the pathological role of the growing cytotoxicity of Aβ pores on glutamate release and the downstream effects on synaptic dysfunction and cognitive decline observed in AD.

## Methods

### Calcium model with Aβ pores

We extend our previous model of CA3-CA1 hippocampal synapses^48^ to incorporate an additional nanodomain, representing Aβ-pore driven Ca^2+^ transients (Fig. 1). This new compartment captures localized Ca^2+^ dynamics produced by spatially distributed clusters of Aβ pores in the plasma membrane, analogous to the VGCC microdomain described extensively in our earlier work^48^. We assume a cytosol-to-pore domain volume ratio of 100 (*δ*_4_), approximating a half-spherical volume of radius 83.5nm, comparable to the Ca^2+^ domain surrounding the IP_3_Rs. As a result of these considerations, the Ca^2+^ dynamics in the respective compartments as well as the pore domain are described by the following equations, while the core model equations and parameters governing the Ca^2+^ fluxes remain unchanged from previous work.

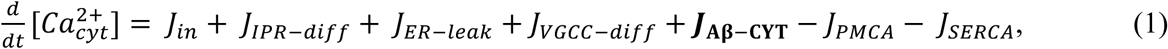

**Figure 1.**
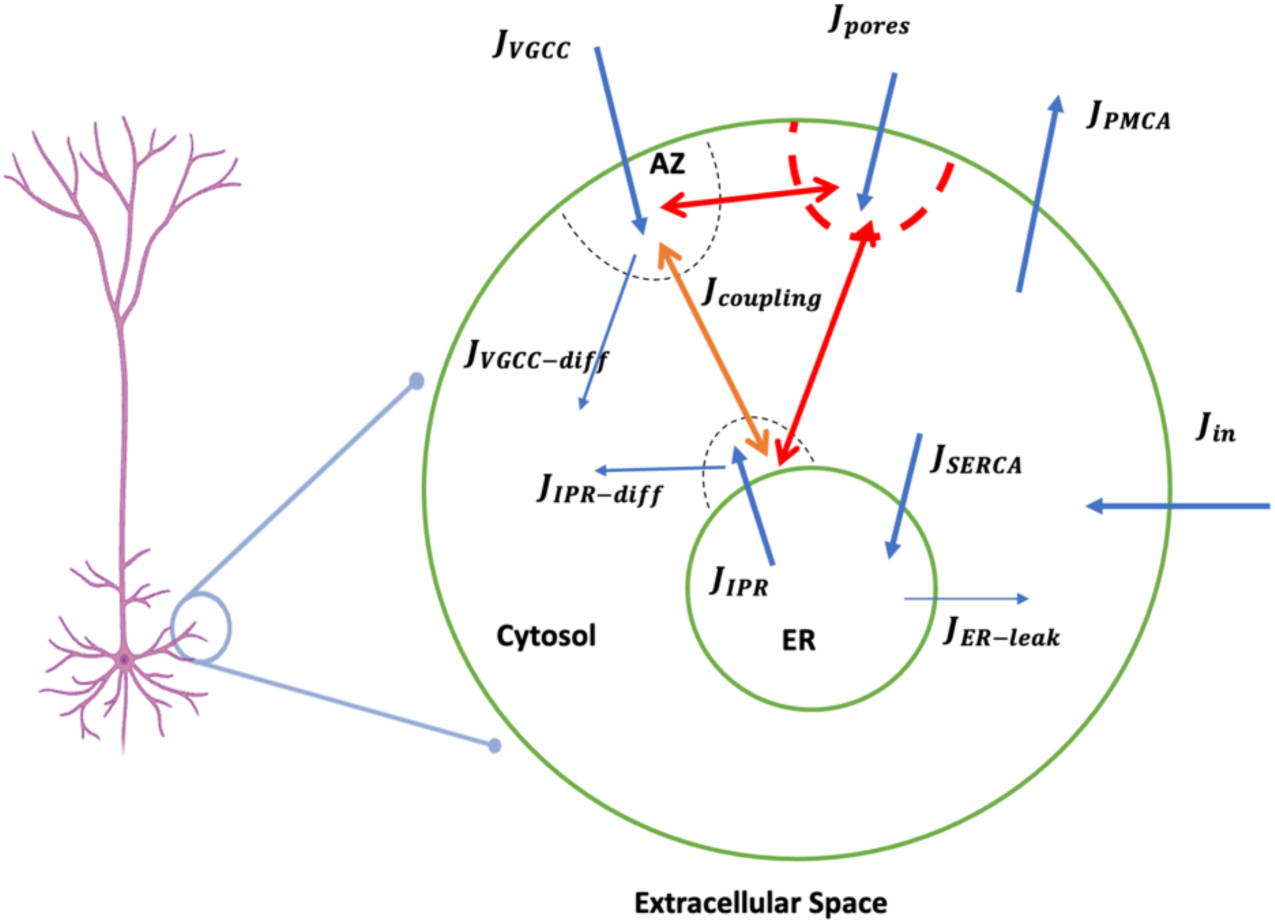
Schematic of the extended multi-compartmental model of Ca^2+^ dynamics in small hippocampal axonal boutons. ^48^: Model representation of the compartmentalized Ca^2+^dynamics, including distinct microdomains surrounding VGCCs, IP₃R, and Aβ pore clusters. The arrowheads represent Ca^2+^ flux between the compartments, and the localized domains are depicted by semicircular boundaries delineating spatiotemporal Ca^2+^ restriction zones.

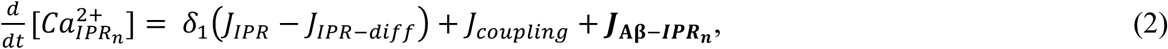

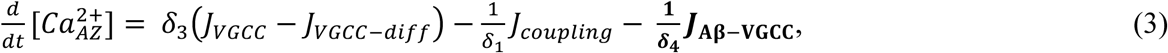

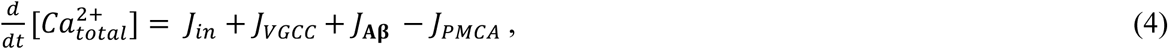

where 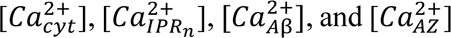 represent the Ca^2+^ concentrations in the cytosol, IP_3_R microdomain, Aβ pore cluster, and VGCCs cluster situated in the AZ, respectively. We modelled the Ca^2+^ influx through Aβ pores (*J*_*Aβ*_) as a passive, electrogenic flux:

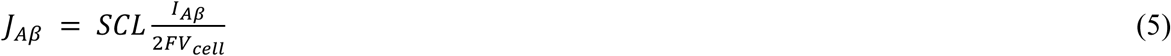

Where *I*_*Aβ* is the total current through all pores in a single cluster, defined as *I*_*Aβ*_ = *pore density* ∗ cluster area_ ∗ *I*_*s*_, with *I*_*s*_ being the single-pore Ca^2+^ current. *F* is the Faraday’s constant, and *V*_*cell*_ is the cell volume. To capture the unconventional gating properties of Aβ pores, we include a sub- conductance/sub-permeability level (*SCL*) parameter which denotes the level at which the pores can gate at a given time and is supported by extensive literature^18,23,38^. We consider SCL values between 1 and 5, in agreement with the previous findings that suggest that Aβ pores display about five distinct peak conductance levels^23^, typically between 0.4 – 4 pS^18^. To account for this, we set the single-pore current to 0.05 pA per permeability level, such that the pore allows a current of 0.05 pA when gating in SCL 1, 0.1 pA when in SCL 2, and so on.

To investigate pore-mediated intracellular Ca^2+^ dysregulation, our model includes additional fluxes invading the bulk cytosol *J*_*Aβ-CYT*_, representing Ca^2+^ diffusion from the Aβ pore nanodomain to the bulk cytosol, modelled as a linear process, and bidirectional coupling fluxes, *J*_*Aβ-IPRn*_ and *J*_*Aβ-VGCC*_, mimicking inter-domain Ca^2+^ exchange in order to capture the effect of pore-driven Ca^2+^ transients on ER Ca^2+^ handling and vesicular fusion at the active zone. Implementing a linear diffusion model becomes numerically intractable at physiologically relevant nanometre scales, hence these coupling fluxes are modelled similarly to *J*_*coupling*_ from our previous work^48^. As a result, *J*_*Aβ-IPRn*_ and *J*_*Aβ-VGCC*_ are defined respectively as,

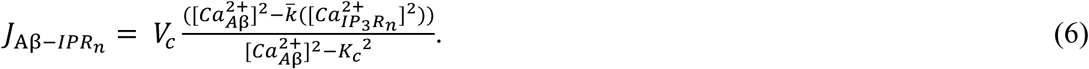

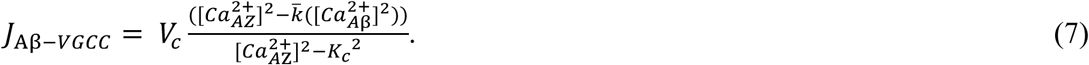

Where, *V*_$_represents the maximum transfer capacity, *K*_*c*_denotes the half-maximal transfer rate, and, *k* indicates the concentrating power of the transfer components, all parameters retain their previously defined values^48^. While this flux representation has limitations, it provides a computationally tractable method to account for the interplay between [*ca*^2+^_*IPRn*_], [*ca*^2+^_*AZ*_], and [*ca*^2+^_*IPRn*_] at physiologically relevant manometer scales, where linear diffusion models become numerically intractable.

### Gating kinetics and temporal evolution of Aβ pores

We previously characterized the dynamics of plasma membrane pores formed during bath application of Aβ_42_ on *Xenopus* oocytes using single-channel imaging^39^. Our findings showed that these pores exhibit increasing Ca^2+^ toxicity over time, characterized by enhanced open probability, larger Ca^2+^ permeability, and a progressive increase in pore population hours following exposure. Notably, after a 25-minute exposure to Aβ_42_ oligomers, the *Xenopus oocyte* expressed on average 16 pores within a 0.3x0.4 μm^2^ membrane patch. Comprehensive statistical analysis of these experiments revealed that these pores exhibit a basal open probability (P_o_) of 0.015, mean open time (τ_o_) of 10 ms, and mean close time (τ_c_) of 2500 ms. To extend our understanding of pore kinetics and evolution beyond the experimental time scales explored in the study, we analyzed single Aβ pore time traces using our proprietary java-based software, TraceSpecks^38,49,50^. Our findings corroborated previous observations in HEK 293 models^23^, indicating that Aβ pores display distinctive gating conductance and time distributions, occasionally manifesting different permeability levels (SCLs) at a given time^23,37,38^.

To simulate the kinetics and temporal evolution of the pores, we adopted the gating properties (*P*_0_ = 0.015, τ_0_ = 10*ms*, and *SCL* = 1) observed after 25 minutes of exposure as the basal gating parameters for our simulations. We modeled the temporal pore behavior within the timescale of our simulations by setting the initial open time to 0 ms and defining the closing time of all pores as the total duration of the simulation. Guided by our previous investigations^39^, we used the mean open duration of the pores, τ_0_, to represent the average temporal extent of a pore opening, while the mean open probability, *P*_0_, quantified the average likelihood of a pore being open at any given moment. To ensure congruence with the previously reported experimental conditions^53,54^, we next estimated the cumulative duration of pore opening by multiplying the mean open probability by the experimental duration, representing the total duration in which the pores remained in an open state. Subsequently, we sampled a series of open times from the available time range, yielding multiple opening events characterized by the specified open and close times. To implement this, we partitioned the available time range into smaller intervals matching the mean open time. From these intervals, we randomly selected open times, representing distinct instances of pore opening events. Each sampled open time effectively denoted a current injection event, enabling us to capture the diverse dynamics exhibited by the Aβ pores. For clarity, we have provided detailed implementation instructions within our MATLAB scripts provided with this manuscript to ensure reproducibility of our simulations. Rate equations were solved using the fourth-order Runge–Kutta algorithm (RK4) with a 1μs time step. All numerical simulations were performed in MATLAB (The MathWorks, Natick, MA, USA).

## Results

### Acute Effects of Aβ Pores on Intracellular Ca^2+^ Signalling and Synaptic Transmission

We first examined the acute effects of pore-driven Ca^2+^ entry on AP-evoked neurotransmitter release and synaptic integrity following a typical 25-minute exposure to Aβ_42_ oligomers. To achieve this, all synaptic configurations were simulated with basal pore gating properties (P_o_ of 0.015, τ_o_ of 10 ms, and τ_c_ of 2500 ms). Additionally, we explored whether the bidirectional coupling between the pore- and IP_3_R Ca^2+^ domains further exacerbates the recently uncovered FAD-driven upregulation of AZ [Ca^2+^] and neurotransmitter release as a result of the gain-of-function enhancement of IP_3_Rs^48^. Our results demonstrate that the presence of Aβ pores dramatically alters exocytosis in response to a single AP (Fig. 2A). Healthy synapses without pores (IP_3_R-WT) display a characteristic sharp, transient release, followed by a rapid return to baseline. In contrast, healthy synapses with pores (Aβ & IP_3_R-WT) exhibit a slightly lower initial peak but a significantly prolonged release phase for over 30 ms, while AD-affected synapses with Aβ pores (Aβ & IP_3_R-AD) displayed an even more pronounced and sustained release profile. AD-affected synapses refer to synapses where IP_3_R exhibits gain-of-function enhancement due to FAD-causing mutations in presenilin. Collectively, these results suggests that although Aβ pores alone can significantly alter the temporal dynamics of exocytosis, the combination of AD-associated IP_3_R dysfunction and Aβ pore formation exacerbates the alterations in exocytosis and leads to a more severe and prolonged disruption of normal synaptic transmission.

**Figure 2.**
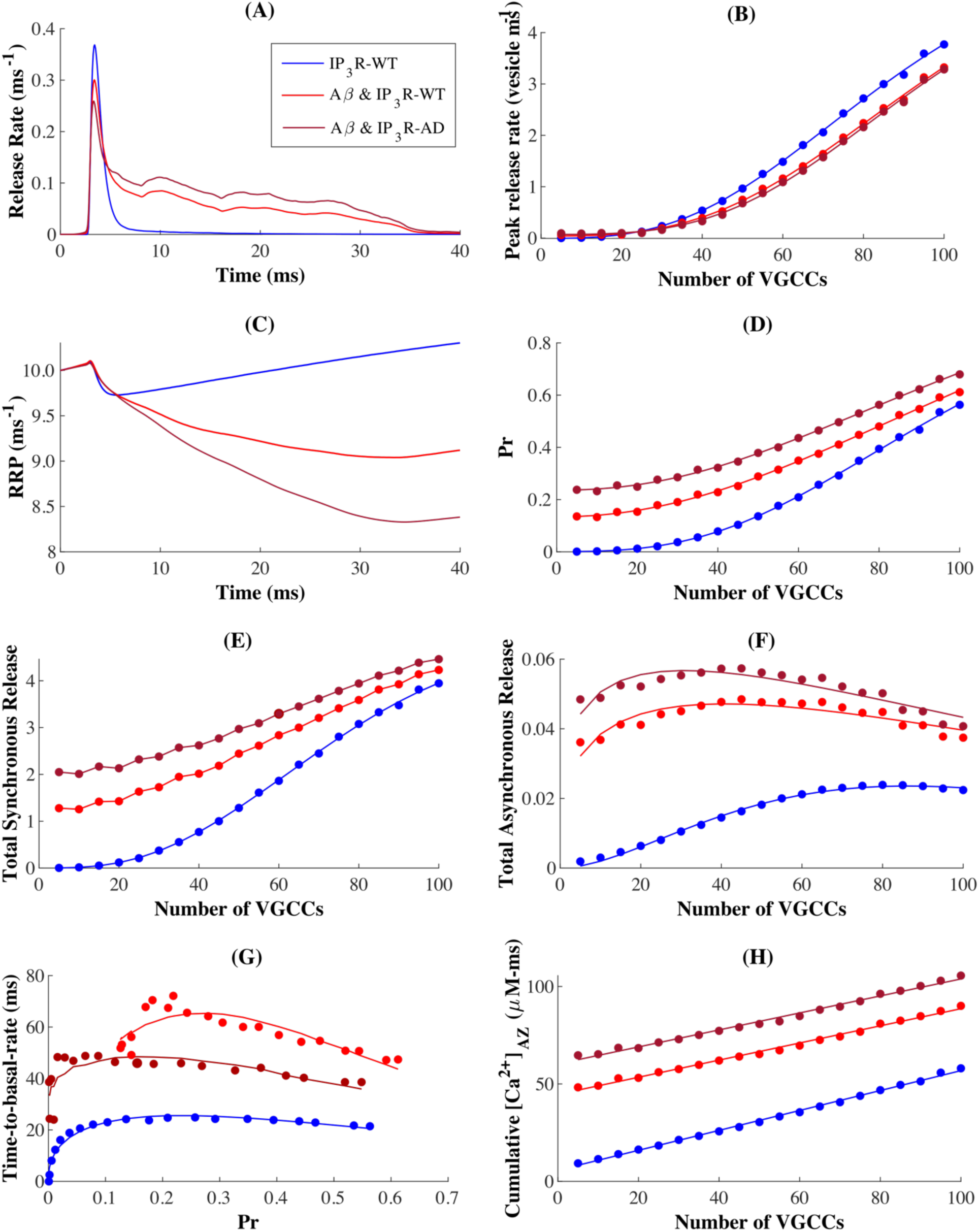
Aβ pore-mediated disruption of synaptic vesicle release dynamics: (A) Neurotransmitter release rate in response to a single AP across IP₃R-WT, Aβ & IP₃R-WT, and Aβ & IP₃R-AD synaptic configurations. (B) Peak release rate as a function of VGCC number. (C) Time course of readily releasable pool (RRP) size following AP stimulation. (D) Release probability (Pr) of synaptic vesicles as a function of VGCC number. (E) Total synchronous release and (F) total asynchronous release modes for varying VGCC distributions. (G) Time-to-basal release rate versus Pr. (H) Cumulative [Ca²⁺]_AZ_ with increasing VGCC expression, demonstrating condition-dependent Ca²⁺ handling.

We then explored whether Aβ pores additionally alter the maximal rate of release rate by exploring how the peak release rate varies with the number of VGCCs (Fig. 2B). Consistent with the characteristic transient peak shown in Fig. 2A, IP_3_R-WT synapses maintained higher peak release rates relative to pore-affected synapses, especially in synaptic configurations containing more Ca^2+^ channels (Fig. 2B). Surprisingly, FAD-driven IP_3_R dysfunction conferred only marginal additional impact on peak release rates in pore-affected synapses, suggesting that the reduced peak release rate is primarily attributable to Aβ pore-driven Ca^2+^ perturbations at the AZ, and operates independently of the intrinsic Ca^2+^ handling machinery. When considered alongside the prolonged release profiles observed in Fig. 2A, this diminished peak release rates in pore-affected conditions reveal that Aβ pores can fundamentally alter the both the temporal dynamics and magnitude of release, where Aβ pores on one hand trigger a broader temporal distribution of release, and on the other lead to persistently diminished release magnitude across a range of VGCC configurations. Given that synaptic vesicle release is primarily coupled to Ca^2+^ in the vicinity of the AZ, these results together provide strong evidence that Aβ-driven pore formation alters the precise spatiotemporal Ca^2+^ signaling necessary for coordinated synaptic transmission regardless of the native Ca^2+^ regulatory mechanisms.

To precisely understand how these pores contribute to impaired vesicle dynamics, we first analyzed the readily releasable pool (RRP) kinetics across the different synaptic configurations (Fig. 2C). IP_3_R-WT displayed an expected pattern of transient RRP depletion followed by gradual recovery shortly after the AP, which reflects the robust vesicle replenishment mechanisms that maintain synaptic efficacy during prolonged periods of activity. However, both synapses with pores exhibited severely altered RRP kinetics, characterized by pronounced and sustained pool depletion that persisted well after the stimulus. Again, this impairment was most pronounced in Aβ & IP_3_R-AD synapses. Considering our model incorporates Ca^2+^-dependent endocytic mechanisms, these results point to an acute failure of such synapses to replenish vesicle stores. In the same vein, given that pool sizes continued to diminish even after cessation of stimulation, the findings here suggest that pore-affected presynaptic terminals may display increased vulnerability to synaptic fatigue during sustained activity. Thus, such perturbations may represent a key mechanism through which Aβ-mediated Ca^2+^ dysregulation may initiate synaptic dysfunction in early-stage AD, particularly through the disruption of vesicle recycling mechanisms.

We next sought to elucidate the functional consequences of Aβ pore-induced perturbations. To achieve this, we assessed the intrinsic probability (P_r_) across a range of VGCC configurations. Our analysis revealed that AD-affected synapses with Aβ pores consistently display the highest P_r_ relative to other synaptic distributions, followed by Aβ & IP_3_R-WT synapses. Such enhancement of P_r_ in Aβ & IP_3_R-AD synapses indicates a reduced threshold for neurotransmitter release that could profoundly increase background noise during stimuli-evoked signaling. We then examined the impact of these alterations on the two key modes of synaptic vesicle release (SVR)— synchronous and asynchronous (Fig. 2E & F). The pattern of total synchronous release (Fig. 2E) closely mirrored the trends observed in release probability, where AD-affected synapses with pores consistently demonstrated markedly increased synchronous release, particularly in configurations with lower VGCC densities. Conversely, IP_3_R-WT synapses maintained the lowest synchronous release across all configurations. Asynchronous release profiles revealed a more complex, non-linear relationship with channel density (Fig. 2F). Both pore-containing synapses—whether AD-affected or healthy—showed elevated asynchronous release compared to control synapses. Interestingly, this effect peaked in synapses with intermediate VGCC distributions, precisely in the range found in small CA3-CA1 synapses in the hippocampus. Given that information encoding in these synapses is tightly coupled to fast synchronous release, this shift towards increased asynchronous release may significantly compromise the temporal precision of synaptic signaling that underlies information processing in hippocampal circuits.

Indeed, vesicle release decay times appear most prolonged in pore-affected synapses. This effect unsurprisingly peaks at intermediate P_r_ values, especially in synapses affected by both FAD and Aβ pores (Fig. 2G), further reinforcing the heightened susceptibility of such small synapses distributions to the temporal perturbations induced by both Aβ pores and AD-associated disruptions. We additionally observe a diminished effect at higher P_r_, indicating that robust Ca^2+^- influx through densely packed VGCCs may partially override any pore-mediated perturbations of intracellular Ca^2+^.

These findings together highlight a previously unrecognized relationship between the intrinsic Ca^2+^-machinery and Aβ pores, revealing that the relative contribution of pore-mediated Ca^2+^ dysregulation to release timing depends critically on the intrinsic release probability of the synapse. To mechanistically explain these temporal alterations, we examined the cumulative Ca^2+^ influx at the active zone (AZ) (Fig. 2H). Indeed, Aβ pores induce persistent elevation of [Ca^2+^]_AZ_, with the most striking effects observed in AD-affected synapses. Thus, the markedly elevated [Ca^2+^]_AZ_ in AD-affected synapses with Aβ pores provides a direct molecular mechanism linking Aβ pore formation to the observed prolonged release events and altered temporal dynamics in affected synapses.

### Pore-driven Ca^2+^ disruptions compromise very short-term plasticity

While it is well established that Aβ forms transient pores that elevate intracellular Ca^2+^ and disrupt synaptic vesicle recycling, the immediate impact of these pores on synaptic plasticity remains unexplored. Thus, we sought to examine how Aβ pore-mediated Ca^2+^ influx influences short-term presynaptic plasticity (STP), using a paired-pulse stimulation protocols outlined in our previous work^48^ (Fig. 3A). Specifically, we probe synaptic response using paired APs delivered at a 40 ms interval to the nerve terminal. The resulting paired-pulse ratio (PPR)—defined as the ratio between second-pulse (Pr_2_) and first-pulse (Pr_1_) release probabilities, averaged across multiple trials— served as the assay of STP. This well-established measure reveals whether synapses undergo short-term facilitation (PPR > 1) or depression (PPR < 1), offering critical insights into rapid synaptic strength modulation.

**Figure 3.**
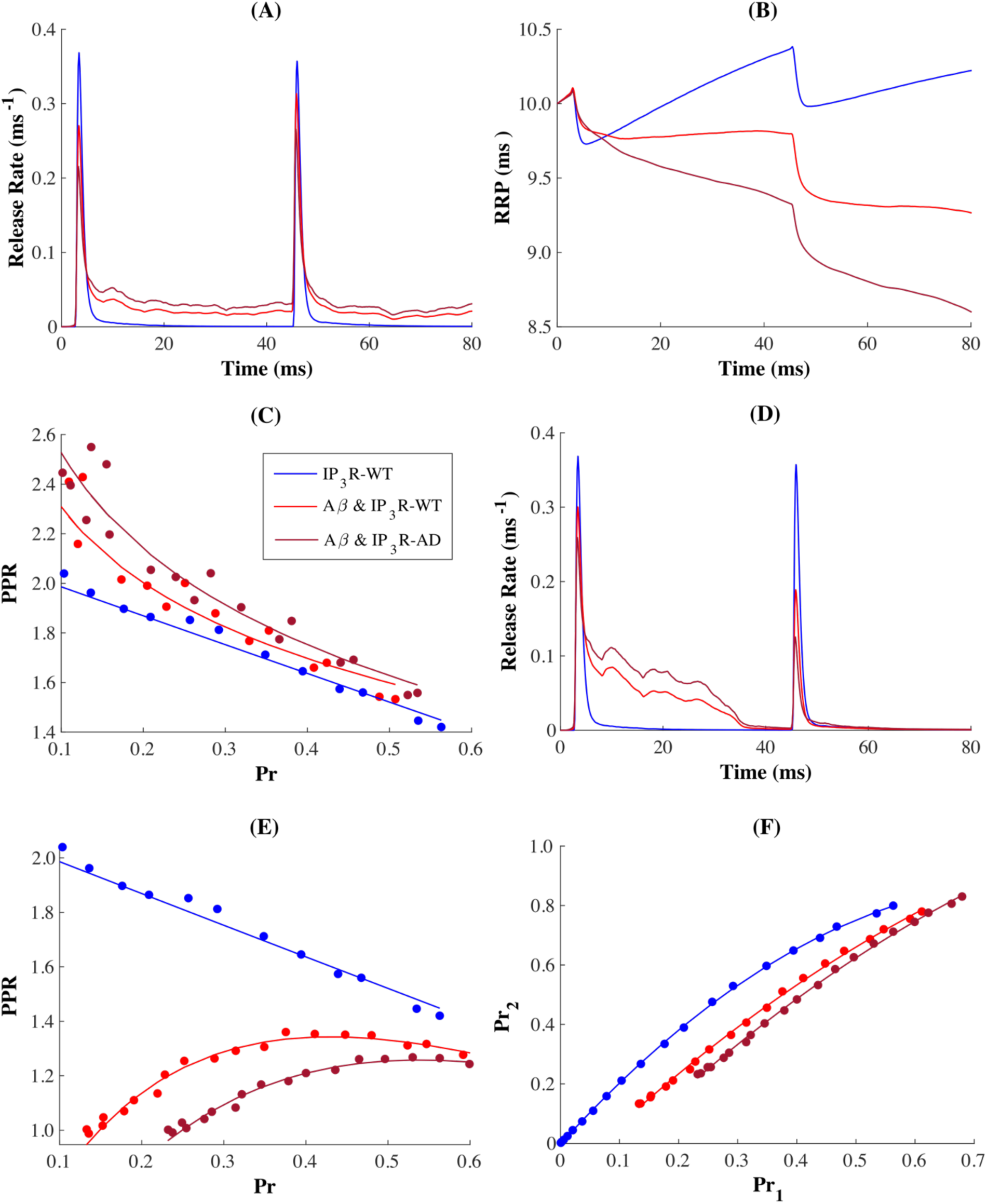
Aβ pores alter short-term plasticity: (A) Release profiles in response to paired-pulse stimulation (40 ms interval) across synaptic configurations. (B) Pore-driven modulation of RRP kinetics. (C) Condition-dependent relationship between PPR and initial release probability (Pr_1_). (D) Temporal evolution of release rates under restricted pore activity, highlighting the phase-specific impact of Aβ pores. (E) Temporal dynamics of pore activation reshape the relationship the dependence PPR on the intrinsic release probability. (F) Release probability in response to the second pulse (Pr_2_) as a function of release probability in response to the first pulse (Pr₁).

Our analysis revealed that Aβ-induced disruptions in release dynamics persist beyond the initial AP, which significantly affects how synapses respond to repeated activation (Fig. 3A).

Specifically, IP_3_R-WT synapses maintained their characteristic temporally precise transient release peaks and rapid recovery for both pulses. In contrast, Aβ & IP_3_R-AD/WT synapses displayed marked alterations, characterized by modest attenuation of initial peaks followed by significantly extended release phases that failed to return to resting value before the second stimulus (Fig. 3A). Most strikingly, the second pulse triggered an even more severe dysregulation of release, pointing to a progressive deterioration of Ca^2+^-dependent vesicle recycling mechanisms (Fig. 3B). The cumulative nature of this impairment reinforces our initial findings that Aβ pores not only disrupt baseline transmission but fundamentally compromise the temporal precision essential for sustained synaptic function during repeated activation. Such activity-dependent perturbations is particularly relevant in hippocampal circuits, where synapses typically operate and respond to high frequency signals^22,23,37,51^.

Further analysis of PPR revealed that all synaptic configurations demonstrated an inverse relationship with the intrinsic response (Pr_1_) (Fig. 3C), reflecting how vesicle depletion constrains subsequent release events, typically superseding any Ca^2+^-dependent facilitation^43,52^ (Fig. 3B). However, both pore-affected synapses displayed distinctive rightward-shifted and steeper PPR-Pr_1_ curves compared to control synapses, indicating that pore formation fundamentally alters the capacity of the synapse to maintain efficacy during repeated activation. Although all configurations demonstrated STF (PPR > 1), the magnitude diminished progressively with increasing Pr_1_, particularly in synapses with more Ca^2+^ channels. This attenuation was most dramatic in synapses affected by both FAD and pores, further suggesting that the convergence of Aβ pores and AD-associated IP_3_R dysfunction generates a particularly severe and unique pattern of release dysregulation. Rather than merely altering baseline responses, together, these pathological changes appear to compress the dynamic range over which synapses can modulate their strength during repetitive activation. Such restriction of STP may help explain how Aβ- mediated Ca^2+^ dysregulation compromises synaptic function—affected synapses lose their ability to encode subtle variations in the timing of incoming stimuli.

To test the veracity of these claims, we performed a targeted set of simulations where pores were only active during the first stimulus of the paired pulse. This approach allowed us to isolate the temporal contributions of Aβ pores to activity-dependent plasticity. Even under restricted activity, pore-affected synapses displayed significant PPR reduction (Fig. 3E), with Aβ & IP_3_R-AD synapses again showing the most pronounced shifts and reduction in PPR. The persistence of these impairments suggests that even brief periods of activity can induce lasting modifications to the capacity for STF in affected synapses. We then examined the role of pores in shaping the relationship between consecutive release events by analysing Pr_2_ as a function of Pr_1_ (Fig. 3F). This analysis represents the conditional probability that a successful release event on first pulse is followed by another successful release on the second pulse. This essentially helps to understand how success in the first release influences the likelihood of success in the second. As with facilitating synapses, our data demonstrate that, across all configurations, the success of a transmission on the second stimulus monotonically depends on release following the first pulse. However, synapses expressing pores activity only during the first pulse displayed notably attenuated facilitation during subsequent activation compared to IP_3_R-WT synapses. These findings reveal an unexpected biphasic role for Aβ pores in modulating synaptic activation. When pores remain continuously active, they promote synaptic hyperactivation. However, temporally restricted pore activation drives synapses to maintain relative hypoactivation over short timescales. This suggests that Aβ pores act as dynamic regulators of synaptic function, with their effects depending strongly on their temporal pattern of their activity. Such nuanced effect may have significant implications for our understanding of pore-driven synaptic dysfunction and could potentially explain some of the varied and heterogeneous synaptic responses to Aβ observed in early-stage AD.

### Temporal Pattern of Aβ Pore Activity Reshapes Synaptic Response During Sustained Activity

To investigate whether these Aβ pore-mediated disruptions extend to sustained synaptic activity, we next examined response profiles during prolonged high-frequency stimulation of 20Hz trains of APs under conditions of unrestricted, where pores remained active throughout the entire stimulation train, and restricted activity, where pore activation was confined to the initial stimulation period. In unrestricted configurations, all conditions showed similar profiles of depression profiles, with responses eventually diminishing to approximately 40% of the initial response by the end of train (Fig. 4A1). Unsurprisingly, the pool of available vesicles showed corresponding pattern of monotonic depletion with consecutive spikes in both control and pore- affected synapses (Fig. 4B1). However, selective restriction of pore activity to just the initial pulse revealed a profound mechanistic vulnerability in Aβ-affected synapses (Fig. 4A2 & B2). Under these conditions, affected synapses showed rapid depression to approximately 20% of their initial response amplitude just within the first five stimuli, contrasting sharply with control synapses that maintained much stronger responses throughout the train. This severe depression corresponds with accelerated RRP depletion in the Aβ-affected (Fig. 4B2), indicating that even brief period of concentrated pore activity can trigger persistent perturbations in the vesicle release mechanisms. Brief periods of intense pore activity thus emerge as critical triggers for the enduring alterations in release machinery, whose effects persist well beyond the temporal window of pore activation. Such temporal sensitivity suggests that early, transient periods of pore formation might be particularly critical in initiating long-lasting synaptic dysfunction, potentially underlying how early Aβ accumulation leads to progressive synaptic deterioration in the early stages of AD.

**Figure 4.**
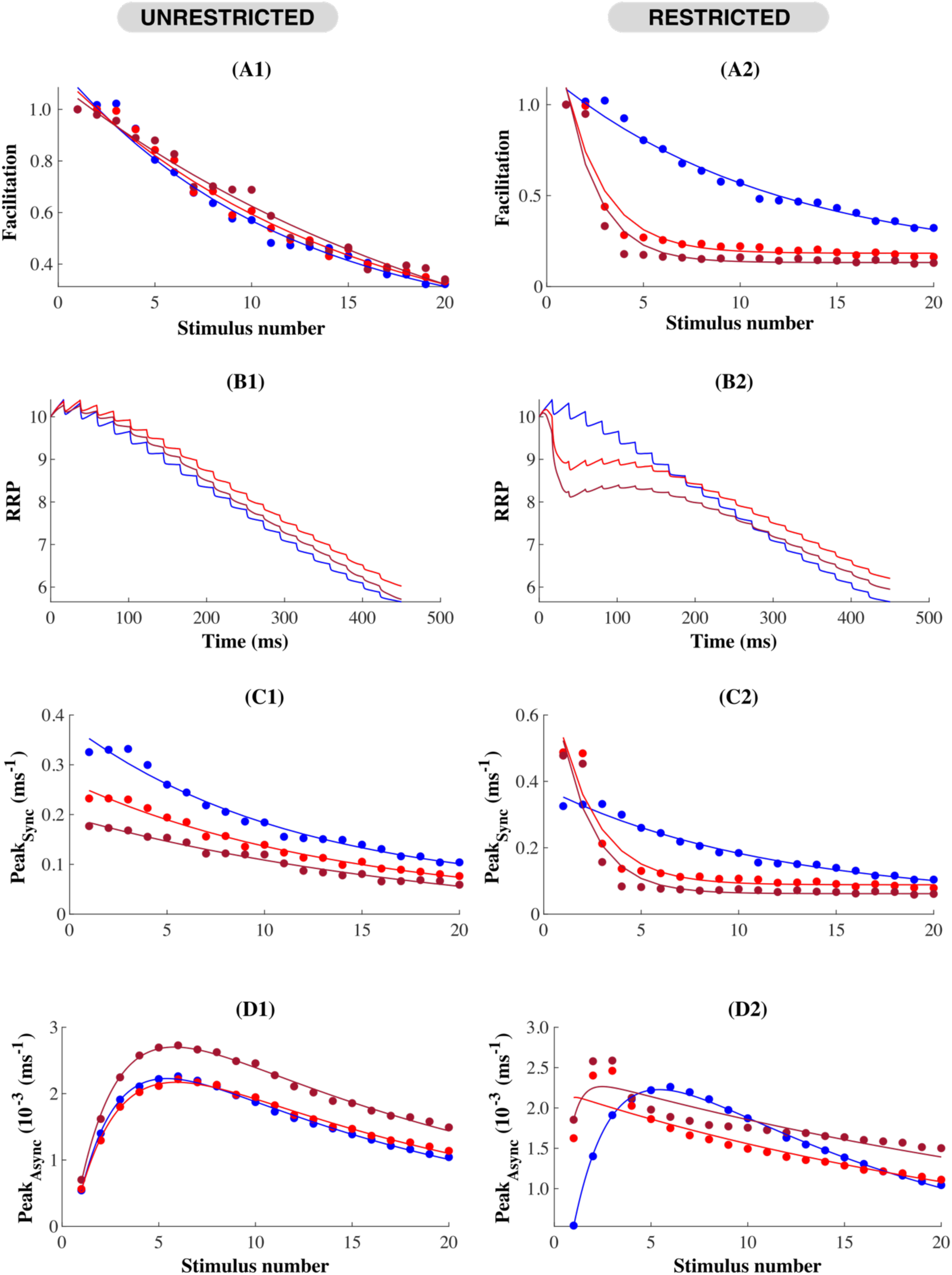
Temporally distinct pore activity differentially regulates synaptic response and vesicle dynamics during sustained activity: (A) Facilitation profile in response to 20 pulse stimulus train delivered at 20Hz. (B) RRP depletion following each AP in the train. Evolution of Synchronous (C) and Asynchronous (D) release peaks during the AP train.

**Figure 5.**
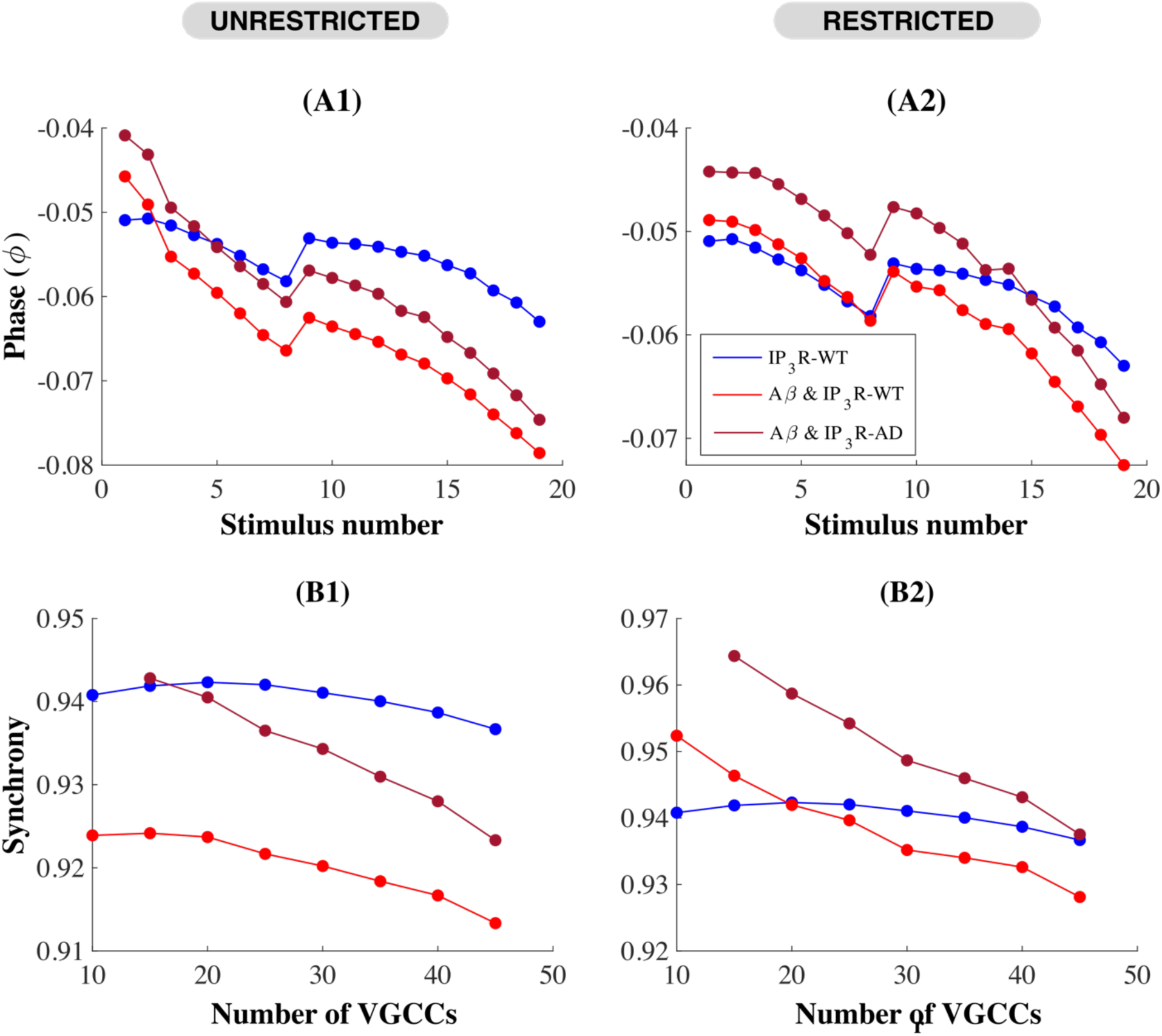
Phase-dependent modulation of synaptic temporal coherence by Aβ pores reveals distinct synchronization patterns in restricted versus unrestricted states: (A) The phase of individual release events with respect to AP in unrestricted (A1) and restricted (A2) pore configurations. Negative phase values indicate release events occurring after the stimulus peak, reflecting the typical delay between stimulus and neurotransmitter release. (B) Overall synchrony of release events in response to stimulus, where higher values along the synchrony axis (approaching 1) indicate tighter temporal coupling between stimulus and release, while lower values suggest more dispersed, less coordinated release timing.

We then examined whether the differential effect of pore-evoked Ca^2+^ dysregulation on release modes (discussed earlier in Fig. 2E & F) persists during ongoing activity. Notably, Aβ-affected synapses exhibited markedly enhanced asynchronous release throughout the stimulus train (Fig. 4D1 & 2), together with significant reduction in synchronous release (Fig. 4C1 & 2). Interestingly, these changes proved independent of the temporal pattern of pore activity, suggesting that Aβ pores fundamentally disrupt the precise spatiotemporal Ca^2+^ signalling that normally maintains the delicate balance between fast synchronous and delayed asynchronous release mechanisms during prolonged activity. While all synaptic configurations displayed the characteristic evolution of the release modes previously described in Ref. ^48^ — marked by progressive decline in synchronous release coupled with biphasic asynchronous release — synapses experiencing restricted pore activation demonstrated distinct temporal signatures. Initial stimuli triggered a pronounced enhancement of asynchronous release followed by acute depletion, with delayed recovery emerging only later in the train. At the same time, synchronous transmission remained persistentlysuppressed, indicating that transient periods of Aβ-mediated Ca^2+^ influx can fundamentally alter the molecular machinery governing how synapses switch between release modes. Mechanistically, the transiently enhanced asynchronous component originates from pathologically elevated residual Ca^2+^ levels, which enable increased competition between lower and higher-affinity Ca^2+^ sensors for vesicles within the RRP. Thus, our findings here together suggest a strong positive correlation between the magnitude of synaptic depression and the transition from synchronous to asynchronous release.

To understand how the pore-mediated disruption of synchronous release affects the temporal fidelity of synaptic transmission, we analysed stimulus-response coordination under both temporally restricted and unrestricted pore activity (Fig. 5). Specifically, we measured event phase vectors (φ, Fig. 5A) and event synchrony (Fig. 5B) following the methods established in previous works^48,53^. Event phase vectors (φ) captured the precise temporal offset between an arriving AP and subsequent neurotransmitter release, measuring stimulus-response coupling at each point in the train. Event synchrony, which was computed as the normalized vector strength of these phase relationships, quantifies how consistently this temporal coordination was maintained across successive stimuli. Together, these metrics provide an assay of the immediate temporal precision of release and the stability of this precision throughout sustained activity. When pores remained active throughout stimulation, all synaptic configurations displayed similar phase patterns for individual release events, characterized by a gradual loss of precision throughout the train, interrupted by a characteristic phase reset in timing after the 9th pulse (Fig. 5A1). However, synapses with only Aβ pores showed significant early-train phase suppression, whereas Aβ & IP_3_R-AD configurations maintained near-normal phase relationships initially before showing marked deviation mid-train. This suggests that Aβ pores alone, without additional pathological effects and under conditions of unrestricted activity, might induce perturbations of release timing more severely than when combined with FAD-associated disruptions, despite apparently normal facilitation profiles.

**Figure 6.**
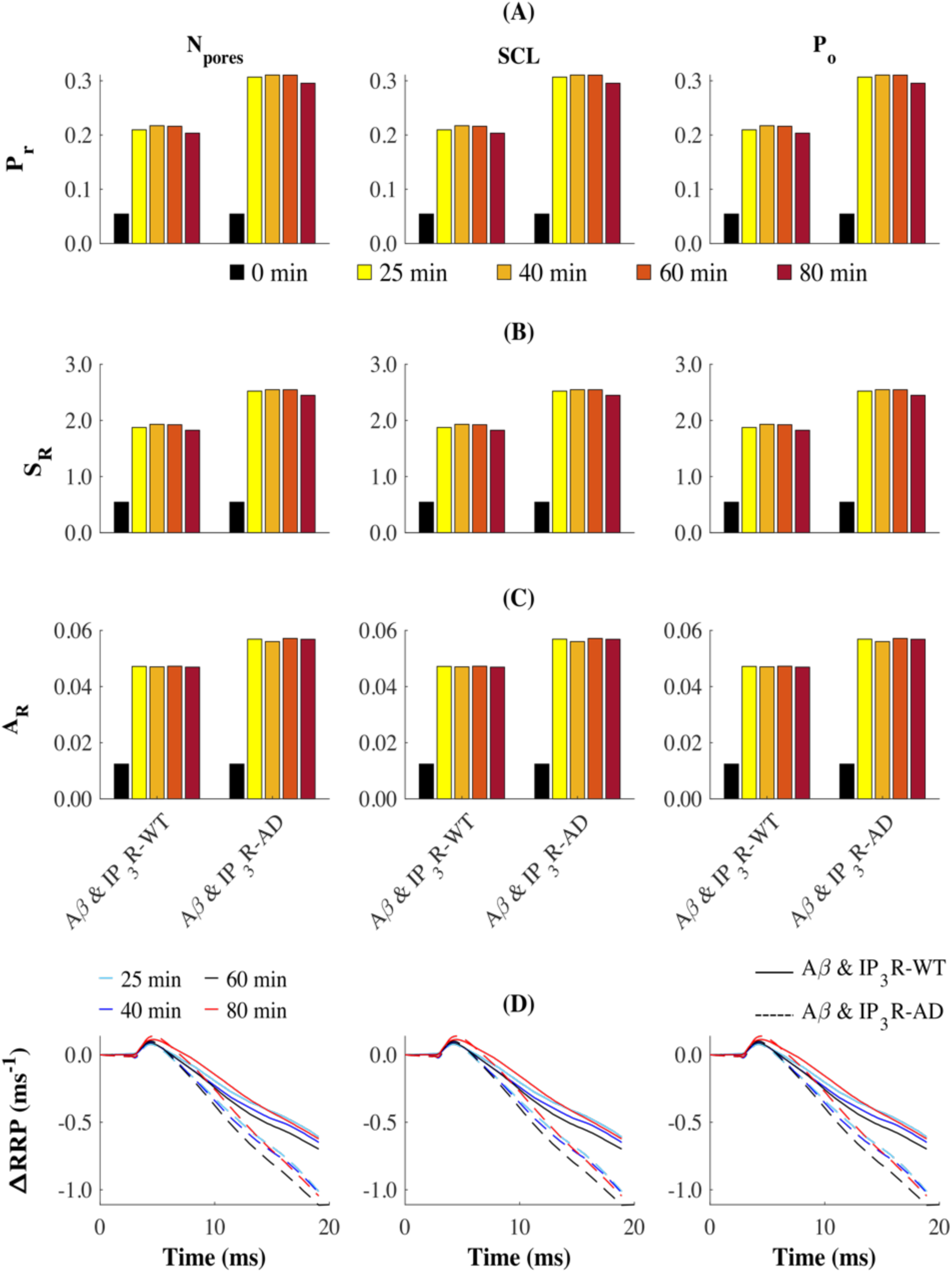
**Temporal evolution of release components during prolonged Aβ_42_ exposure**. Pr (A), synchronous release (S_R_) (B), and asynchronous release (A_R_) (C), across different exposure durations (25-80 min) for varying pore parameters: number of pores (N_pores_, left), sub-conductance levels (SCL, middle), and open probability (Po, right) in both pore-affected wild-type (Aβ & IP_3_R- WT) and AD-affected (Aβ & IP_3_R-AD) synapses. Black bars indicate baseline measurements at 0 min. (B) S_R_ showing temporal progression under increasing exposure durations for each pore parameter. (C) A_R_ remain stable despite prolonged exposure across all pore configurations. (D) Time course of synaptic depression across exposure durations.

Most surprisingly, temporal restriction of pore activity revealed an unexpected enhancement in stimulus-response phase coherence in pore-affected synapses compared to both IP_3_R-WT configurations early in the train, followed by progressive decline (Fig. 5A2). We propose that this biphasic behaviour, on one hand, could be mechanistically explained by the initial rapid asynchronous release triggered by concentrated pore activity (Fig. 4D2). This could subsequently promote compensatory enhancement of more temporally precise synchronous exocytosis (Fig. 4C2). On the other hand, however, broad, unrestricted pore activity leads to sustained elevation of background Ca^2+^ in the AZ driving pronounced asynchronous release and consequently degrading the stimulus-response coordination (Fig. 4D1). Overall response synchrony further revealed that the timing of pore activity differentially alters temporal coordination of synaptic activity. Under continuous pore activity, affected synapses displayed markedly reduced synchrony relative to IP_3_R-WT synapses. However, FAD-affected synapses demonstrated enhanced stimulus-response concordance under temporally constrained pore activation, particularly at physiological VGCC densities, before converging with control values at higher channel numbers. This precise dependence on the activation pattern of the pores demonstrates that the temporal characteristics of pore formation critically determine how Aβ disrupts synaptic information encoding.

### Extended Pore Activity Produces Limited Additional Disruption of Release Machinery Beyond Initial Pore Formation

Previous work has demonstrated that pores formed due to Aβ_42_ oligomers become increasingly active and permeable to Ca^2+^ during extended exposure^38,39^. We reasoned that we could examine how this progressive toxicity affects exocytotic machinery by modelling escalating pore-mediated Ca^2+^ disruption in synaptic configurations with 35 VGCCs. To achieve this, we simulated extended pore exposures to Aβ_42_ oligomers at 40, 60, and 80 minutes, by progressively doubling three key gating parameters at each timepoint: the number of pores (N_pores_), their conductance (SCL), and their open probability (P_o_) during a single AP stimulation (Fig. 5). Our results surprisingly show that extended exposure beyond the initial 25-minute period produces only modest additional alterations in the release components (Fig. 5). Specifically, the elevated release probability (P_r_) and synchronous release (S_R_) seen shortly after initial pore formation showed only marginal further increases between 25 and 60 minutes across all gating properties (Fig. 5A). This was followed by a subtle but consistent reduction at 80 minutes, suggesting potential deterioration of the release machinery. However, asynchronous release (A_R_) remained notably stable throughout extended exposure periods, showing neither the transient elevation observed in other components nor late-phase reduction at 80 minutes (Fig. 5C). The evolution of synaptic depression further supports these observations since the depression profiles remain comparable across the different exposure durations (Fig. 5D). Taken together, these findings suggest that while prolonged exposure to Aβ_42_ and sustained pore formation may enhance overall cellular damage, their direct effects on basic release machinery appear to saturate early in the pathological cascade. This rapid stabilization despite growing molecular disruption may indicate engagement of compensatory or homeostatic mechanisms that initially buffer against escalating Ca^2+^ dysregulation. This suggests that the initial pore formation, rather than their extended activity, may represent the critical trigger for subsequent synaptic dysfunction.

## Discussion

The pathological cascade leading to synaptic dysfunction in AD has remained a subject of intense investigation, with mounting evidence suggesting that soluble Aβ oligomers play a central role through disruption of Ca^2+^ homeostasis^13–15,18^. While numerous studies have demonstrated that Aβ oligomers can form Ca^2+^-permeable pores, the precise mechanisms and spatiotemporal consequences of such perforations on synaptic function have been difficult to assess due to technical limitations in simultaneously monitoring multiple subcellular Ca^2+^ domains^18,23,37,39^. Here we present an extension of our previously proposed multicompartmental model^48^ that incorporates a spatially resolved microdomain to capture transient pore-driven Ca^2+^ influx. This enhanced framework allows us to dissect how the evolving Aβ pore activity interfaces with the endogenous Ca^2+^-handling machinery at presynaptic terminals. Overall, by integrating experimental data on pore kinetics with established mechanisms of presynaptic Ca^2+^ handling, our work here uncovers multiple distinct mechanistic pathways through which Aβ pore formation may drive the early and progressive deterioration of synaptic function characteristic of AD.

Our results demonstrate that Aβ pores fundamentally disrupt the pristine spatiotemporal organization of Ca²⁺ signalling that orchestrates neurotransmitter release. Specifically, we observe that rather than simply elevating or suppressing synaptic transmission, Aβ pores profoundly reorganize the temporal dynamics of vesicle release, which consequently affects the precision and reliability of synaptic response. Indeed, earlier studies have attributed pore-mediated elevation of intracellular Ca²⁺ to either an enhanced or diminished synaptic transmission in AD models^23,37^, however, these apparently contradictory findings have no unified mechanistic explanation. Our work helps resolve this paradox by demonstrating that the effects of these pores depend critically on both their pattern of activity and the properties of the affected synapse. We found that pore formation induces a complex phenotype characterized by reduced peak release amplitude and pathologically sustained neurotransmitter release, with the effect manifesting most dramatically in synapses with physiologically relevant VGCC distributions. This selective disruption becomes especially apparent across the distinct temporal domains of synaptic function. Aβ-affected synapses exhibit enhanced release during brief stimuli but show marked depression during sustained activity. This biphasic behaviour is likewise most striking in synapses operating at low and intermediate release probabilities, a configuration that typically allows hippocampal synapses to maintain reliable transmission despite their inherent variability. In these synapses, Aβ pores appear to drive a transition from tightly coordinated to temporally dispersed release events, which could consequently alter how temporal information is encoded. These perturbations are further exacerbated when Aβ pores act in concert with FAD-associated pathology. The findings here suggest that early synaptic dysfunction in AD, which has traditionally been attributed to gross disruption of Ca²⁺ homeostasis, may instead arise from subtle perturbations of the finely-tuned spatiotemporal Ca²⁺ signals that normally ensure synchronized neurotransmitter release. This distinction reveals why deficits in synaptic response timing can emerge before overt Ca²⁺ dysregulation and provides a mechanistic framework for understanding the progressive deterioration of circuit function in AD, particularly in networks requiring precise temporal coding. Moreover, the time-dependent nature of these effects may help explain the selective vulnerability of specific neural circuits and suggests new therapeutic directions focused on restoring temporal coordination rather than simply normalizing bulk Ca²⁺ levels.

Previous investigations of the neurotoxic effects of these pores have indeed documented their remarkably complex temporal evolution, yet the functional consequences of their unstable conductance and atypical gating properties during sustained activity remain largely unexplored. Our findings here address this question and reveal that these dynamic channel properties could critically shape pore-mediated neurotoxicity in ways not previously recognized. We specifically examine phase-dependent pore activity and uncovered distinct temporal windows where Aβ pores differentially impact synaptic function. For instance, during paired-pulse protocols, we observed that continuous pore activity promotes synaptic hyperactivation, while temporally restricted activation leads to a distinct pattern of hypoactivation at short timescales. Likewise, transient pore activation over longer timescales following a train of AP induces dramatic synaptic depression and accelerated vesicle pool depletion, contrasting with the relatively normal depression profiles seen with unrestricted pore activity. Perhaps most strikingly, temporally brief periods of pore activity transiently enhance early stimulus-response phase coherence before leading to progressive decline. On the other hand, continuous pore activity consistently degrades synchrony, suggesting that the temporal characteristics of pore formation critically determine how Aβ disrupts synaptic information encoding in early AD pathogenesis. Together, these findings suggests that even brief periods of pore formation can trigger persistent alterations in presynaptic release machinery and challenges the traditional view that continuous pore activity is necessary for synaptic dysfunction.

Here, we propose that rather than acting as parallel pathological processes, Aβ mediated membrane perforations and ER Ca²⁺ disruptions function as an integrated pathological unit through bidirectional coupling between their microdomains. While both pathological mechanisms originate via distinct signalling cascades^16,54^, mounting evidence suggests that their convergence on Ca²⁺ dyshomeostasis could work synergistically in both sporadic and FAD^55^, particularly given that FAD mutations can lead to enhanced production of aggregation-prone Aβ_42_ oligomers^56–58^. Our model demonstrates that the combined impact of these perturbations creates multiple feedback loops that amplify the initial disruptions, producing both additive and non-additive effects across the different temporal domains of synaptic function. Specifically, synapses affected by both Aβ pores and FAD-driven IP_3_R dysfunction exhibit fundamentally altered response profiles during single AP stimulation, characterized by elevated Ca²⁺ concentrations in the AZ and prolonged release events. This effect becomes even more pronounced during paired-pulse stimulation, where FAD-affected synapses with pores demonstrate markedly enhanced facilitation compared to synapses affected by either mechanism alone. This enhanced facilitation appears mechanistically lined to the bidirectional coupling between pore-driven Ca²⁺ influx and IP_3_R-mediated Ca²⁺ release, which establishes feed-forward loops that amplify local Ca²⁺ concentrations at the AZ. However, this mutual potentiation becomes detrimental during sustained high-frequency stimulation, where the combined effects trigger severe RRP depletion and profound synaptic depression. The observation that Aβ pores, while developing after the onset of FAD pathology, shows such profound interaction with ER dysfunction may help explain why therapeutic strategies targeting either pathway alone show limited efficacy. These findings suggest that effective AD treatments may need to simultaneous target both membrane integrity and ER Ca²⁺ handling to successfully prevent progressive synaptic dysfunction.

Although Aβ pores have been traditionally viewed as driving ever-increasing neurotoxicity^23,37,39^, our model here reveals a more nuanced temporal evolution of their effects on synaptic function. Indeed, we^39^ and others^23,37^ have documented that these pores exhibit growing toxicity during initial exposure, particularly within the first 30 minutes^23,37^, the consequences of prolonged exposure remains unclear. We therefore address the critical question of whether the progressive synaptotoxicity driven by Ca²⁺ dysregulation continues to increase monotonically with the duration of Aβ exposure. Our results show that while the initial 25–60-minute window shows progressive enhancement of Ca²⁺ influx and synaptic disruption, these effects may eventually reach a plateau rather than escalating indefinitely. This stabilization appears to result from complex interactions between Ca²⁺-dependent mechanisms that ultimately establish a new, albeit pathological, steady state. During this later phase, the impact of Aβ pores on bulk cytosolic Ca²⁺ likely becomes partially decoupled from their effects on submembrane Ca²⁺ domains, especially without additional ER Ca²⁺ mobilization. This stands in contrast to previous studies that focused primarily on early-phase effects or used simpler models of Ca²⁺ dynamics^59^. More so, this temporal evolution manifests differently across various aspects of synaptic vesicle release. Asynchronous release quickly reaches saturation shortly after exposure, while the release probability and RRP dynamics continue evolving at a slower rate. These distinct profiles suggest that initial Aβ-mediated disruptions may trigger compensatory mechanisms that partially stabilize Ca²⁺ homeostasis, even as broader synaptic dysfunction persists. Importantly, this final plateau state is not uniform across all synapses but depends critically on their initial synaptic properties and the presence of FAD-associated pathology, with some synapses showing greater capacity for compensatory adaptation than others. These observations have particularly important therapeutic implications strategies targeting prolonged Aβ aggregation. Interventions may need distinct strategies for the acute phase of rapidly escalating disruption versus the later phase of sustained but stable dysfunction. Moreover, this biphasic progression helps reconcile apparently contradictory reports regarding the effect of Aβ on neuronal excitability, suggesting that observed differences may partly reflect examination at different temporal stages of the pathology.

We note that while our study provides critical insights into how Aβ-pores progressively disrupt synaptic function, there are several important limitations that warrant consideration. Though the current framework captures the complex interaction between Aβ pores and ER Ca²⁺handling, it necessarily simplifies several aspects of synaptic physiology to remain computationally tractable. The compartmentalized approach to Ca²⁺ dynamic, while effective, may not fully capture the complex spatial organization of Ca²⁺ microdomains in real synapses. Similarly, our representation of Aβ pore clusters assumes uniform cluster distribution across the membrane, whereas current experimental evidence suggests they may cluster uniquely in specific membrane regions. *in vivo* effects likely involve complex network interactions that could amplify or attenuate the phenomena we describe. Hence, our focus on single synapses in isolation could preclude inference of network-level effects that may be crucial for understanding how local synaptic dysfunction translates to circuit-level disruption in AD. While our predictions about the plateauing of pore effects are mechanistically plausible, technical limitations in maintaining stable recordings beyond 60 minutes make direct experimental validation challenging. Additionally, our model does not account for potential contributions from other Ca²⁺ -handling organelles, particularly mitochondria, which are known to be early targets in AD pathogenesis and may significantly influence the temporal evolution of Ca²⁺ dysregulation. Despite these limitations, the model successfully reproduces key observations about Ca²⁺ dynamics and synaptic function in both healthy and AD-affected synapses, suggesting that the simplified framework captures essential aspects of the underlying biology. Therefore, while not diminishing the core insights of our study, these limitations point to important future research directions, including the need for improved experimental techniques to monitor pore dynamics over extended periods and more sophisticated computational approaches that can incorporate additional levels of biological complexity and dysfunction in AD pathogenesis.

## Author Contributions

Conceptualization, T.A. and G.U.; methodology, T.A.; validation, T.A. and G.U.; formal analysis, T.A.; investigation, T.A. and G.U.; resources, G.U.; data curation, T.A.; writing—original draft preparation, T.A.; writing—review and editing, G.U.; visualization, T.A.; supervision, G.U.; funding acquisition G.U. All authors have read and agreed to the published version of the manuscript.

## Data Availability Statement

The complete model code as well as analysis scripts will be posted on our lab’s webpage after the manuscript is accepted for publication.

